# Venoms of related mammal-eating species of taipans (*Oxyuranus*) and brown snakes (*Pseudonaja*) differ in composition of toxins involved in mammal poisoning

**DOI:** 10.1101/378141

**Authors:** Jure Skejic, David L. Steer, Nathan Dunstan, Wayne C. Hodgson

## Abstract

**Background:** Taipans of the genus *Oxyuranus* are predominately mammal-eating specialists and a majority of Australian brown snakes of the sister genus *Pseudonaja* are generalist predators, feeding on mammals, lizards and frogs. In this paper, venom composition of several related mammal-eating species was compared using shotgun proteomics.

**Results:** Venom of *Oxyuranus temporalis* consisted predominately of α-neurotoxins (three-finger toxin family) and was deficient in phospholipase A_2_ neurotoxins. In contrast, PLA_2_ neurotoxins (taipoxin and paradoxin) were abundant in the venoms of other mammal-eating taipan species – *Oxyuranus scutellatus* and *O. microlepidotus*. Variation in neurotoxic PLA_2_ expression was also recorded in mammal-eating brown snakes, some species having high venom levels of textilotoxin or related homologues, for example *Pseudonaja textilis* and *P. nuchalis*, and others, such as *P. ingrami*, lacking them. Venom prothrombinase proteins (fX and fV) were expressed in most mammalivorous lineages, being particularly abundant in some *Pseudonaja* species. Notably, *Oxyuranus temporalis* venom was deficient in venom prothrombinase despite a mammal-based diet. Expression of an α-neurotoxin that is lethal to rodents (pseudonajatoxin b) was profoundly down-regulated in *Pseudonaja textilis* venom sample from Queensland and highly up-regulated in the sample from South Australia despite a report that the snake feeds on rodents in both regions.

**Conclusion:** Related species of taipans and brown snakes that feed on small mammals express different sets of venom proteins toxic to this vertebrate group. This suggests an involvement of factors other than prey type selection in shaping venom proteome composition.

## Introduction

Variation in snake venom composition has attracted a lot of research interest, but the bases of this phenomenon remain poorly known. A recent review of snake venom proteome profiles emphasised the presence of familial trends in venom composition – venoms of viperid snakes generally abound in metalloproteinases, phospholipases A_2_ and serine proteases, and those of elapids are dominated by three-finger toxins and phospholipases A_2_ [1]. However, marked intrafamilial variation is also present. For instance, venom of the black mamba (*Dendroaspis polylepis*), an African elapid that preys on mammals, is predominately composed of kunitz-type dendrotoxins and 3FTxs, whereas the venom of a distantly related Australian mammalivorous elapid, the coastal taipan (*Oxyuranus scutellatus*), contains PLA_2_s and venom prothrombinase [1-3]. Types and quantities of secreted venom proteins often vary among closely related species and populations. Protein composition differences in venoms among related species can be very large, for example in American coral snakes (*Micrurus*), palm pitvipers (*Bothriechis*) and African Gaboon and puff adders (*Bitis*) [4-6]. In other cases, there seems to be little intragroup variation, for instance in North American pitvipers of the genus *Agkistrodon* [7]. Variation in venom composition is commonly observed between conspecific populations. A well-known example is the Mojave rattlesnake (*Crotalus scutulatus*), which exhibits extraordinarily large differences in the venom content of crotamine-like small myotoxins, metalloproteinases, and PLA_2_s across populations [8].

The causes of variation in venom composition have been hypothesised to result from factors such as diet, phylogeny and geographic distance. A study that examined the effects of these factors on the venom composition of Malayan pitviper (*Calosellasma rhodostoma*) populations concluded that the intraspecific variation in venom composition was closely associated with diet and rejected the effects of geographic proximity and population phylogeny [9]. Another study conducted on North American rattlesnakes of the genus *Sistrurus* found that interspecific variation in abundance of some venom proteins was related to diet [10]. Yet other studies on tiger snake (*Notechis*) populations in South Australia or carpet vipers (*Echis*) found no obvious link between venom composition and the kind of prey consumed [11, 12]. Research on neotropical palm pitvipers (*Bothriechis*) revealed large interspecific differences in venom proteome composition despite similar generalist diets [6, 13].

Australian brown snakes (*Pseudonaja*) and taipans (*Oxyuranus*) are swift, mostly medium to large-sized terrestrial snakes, which form a monophyletic group among the oxyuranine elapids [14]. They are responsible for a number of serious and fatal envenomation cases [15-17]. Taipans are mainly specialist predators of mammals [18, 19]. On the other hand, brown snakes are generalist predators as adults, feeding on mammals, lizards, and frogs [20]. Shine recovered only rats from preserved *P. ingrami* [20], but based on anecdotal reports the species may feed on frogs as well (N. Dunstan, personal communication). *P. modesta*, a small-bodied species, is saurophagous i.e. preying predominately on lizards [20].

Venoms of these elapids contain proteins belonging to several protein families [21-23]. This project focused on expression patterns of proteins that have been shown experimentally to be toxic to rodents and are presumed to have a role in mammal prey incapacitation. These include procoagulant venom prothrombinases and neurotoxins from the phospholipase A_2_ and three-finger toxin families. Venom prothrombinase is a protein complex consisting of coagulation factor X-related serine protease and coagulation factor V-related multicopper oxidase [24-27]. Functionally, it is one type of prothrombin-activator [28]. Upon injection in rodents, this toxin induces coagulopathy and collapse of the circulatory system, manifested in a rapid fall in systemic blood pressure [29]. Neurotoxins target the neuromuscular system and induce paralysis. They belong to two major protein families: phospholipases A_2_ (PLA_2_) and three-finger toxins (3FTX). Not all PLA_2_s are neurotoxic, but some induce potent presynaptic neurotoxicity and are called β-neurotoxins [30, 31]. Neurotoxic PLA_2_s from Australian elapid venoms include textilotoxin, taipoxin and paradoxin and display variable protein complex composition [32-34]. 3FTxs are a large protein family that includes members with diverse functions. Those that target acetylcholine nicotinic receptors are called postsynaptic or α-neurotoxins [35, 36]. Within this family there are three major clades of α-neurotoxins, including: short-chain (also called type I), long-chain (type II) and oxyuranine (type III) [37, 38].

The aim of this study was to gain insight into venom composition of related mammal-eating species of Australian elapid snakes (Fig.1). Venom composition was analysed using bottom-up shotgun proteomics and relative label-free quantification (iBAQ). Toxin expression profiles and prey types were plotted onto the species phylogenetic tree.

**Figure 1.**
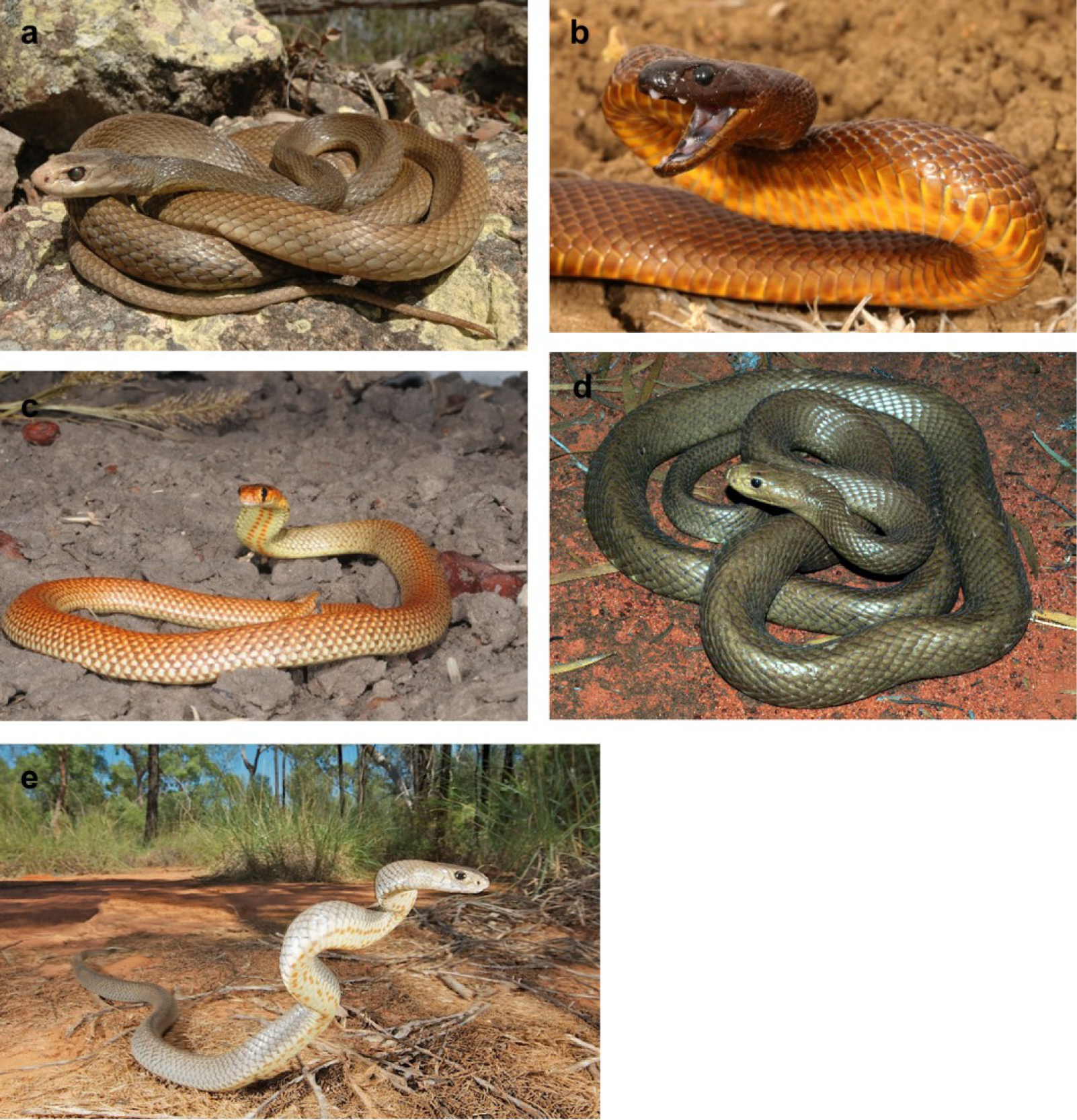
Australian taipans and brown snakes represent an attractive model system to study variation in venom composition. Related mammal-eating specialist and generalist species secrete venoms that differ in the composition of mammal-targeting toxins: (a) coastal taipan (*Oxyuranus scutellatus*), (b) inland taipan (*O. microlepidotus*), (c) Ingram’s brown snake (*Pseudonaja ingrami*), (d) Western Desert taipan (*O. temporalis*) and (e) eastern brown snake (*P. textilis*). Photos taken by Scott Eipper (a and c), Akash Samuel (b), Brian Bush (d) and Stewart Macdonald (e).

## Results and Discussion

### Venom of mammal-eating Western Desert taipan consists predominately of three-finger α-neurotoxins and is deficient in PLA_2_ neurotoxins and venom prothrombinase

Alpha-neurotoxins comprised most of the venom proteome of the Western Desert taipan (*Oxyuranus temporalis*) (Fig. 2). This finding confirms that the very rapid neurotoxic activity of *O. temporalis* venom in chick biventer cervicis physiological assay, reported by Barber et al [39], is due to an extremely high concentration of postsynaptic α-neurotoxins (iBAQ abundance = 97.9%). A recent study sequenced the α-Elapitoxin-Ot1a short-chain α-neurotoxin from this species and found it to be a potent and abundant venom constituent [40]. Chains forming presynaptically-neurotoxic phospholipase A_2_ protein complexes were lacking, with the exception of one protein match that is homologous to the paradoxin alpha chain (Additional files 1-3). The expression level of this chain was so low (iBAQ = 0.28%) that it is unlikely to play a major role in venom neurotoxicity. A very important finding is a deficiency of venom prothrombinase in *O. temporalis* venom. Specifically, a MaxQuant search failed to detect tryptic peptides matching venom fX serine protease in this venom sample, indicating that the enzyme is either extremely scarce or absent. Four peptides corresponding to fV were detected in *O. temporalis* venom, but the overall expression of the protein was negligible (iBAQ = 0.00033%) (Additional files 1-3). Relative abundances of venom protein families for this and other species are listed in Table 1.

**Table 1.**
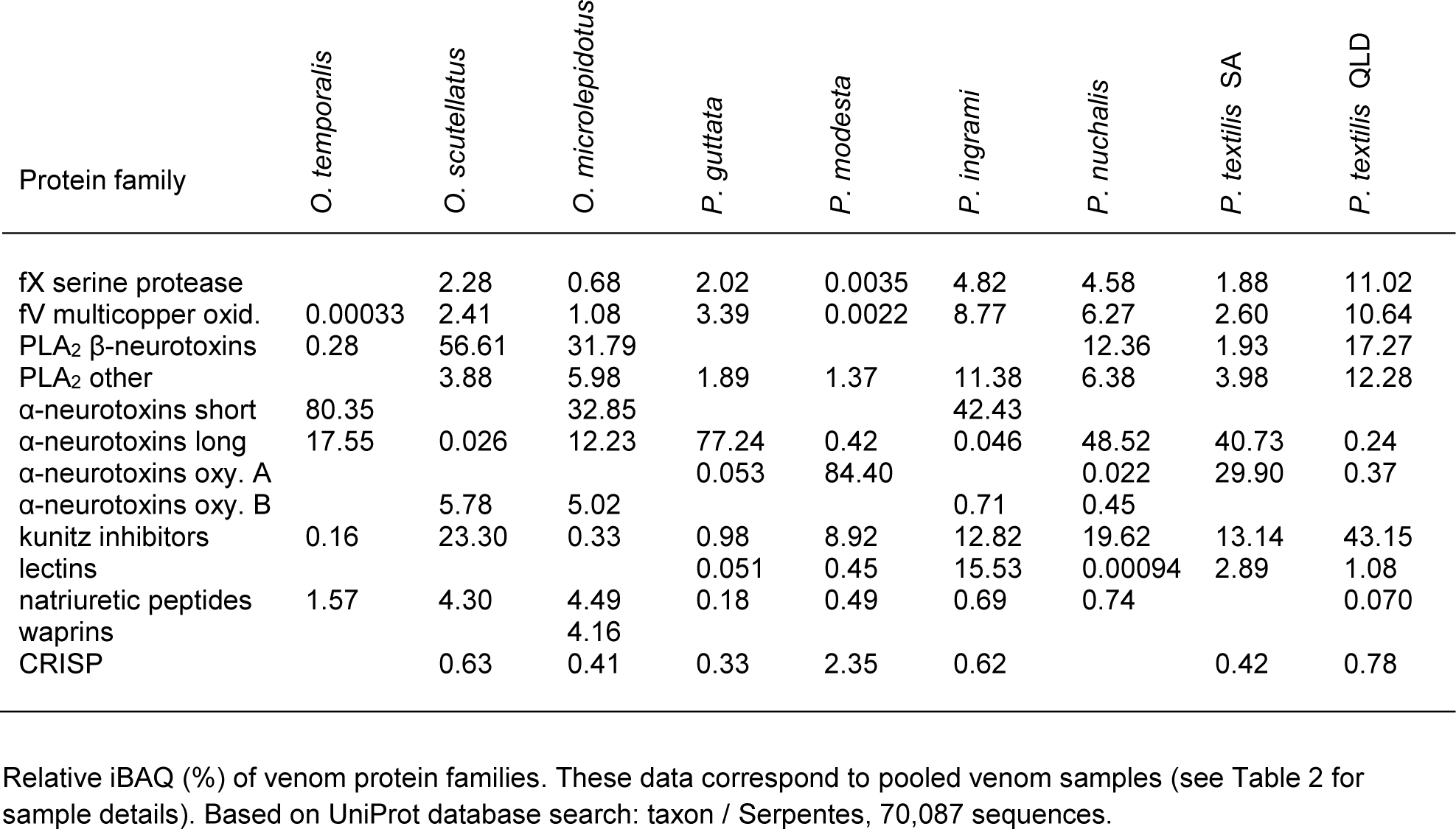
Expression levels of major venom protein families

**Figure 2.**
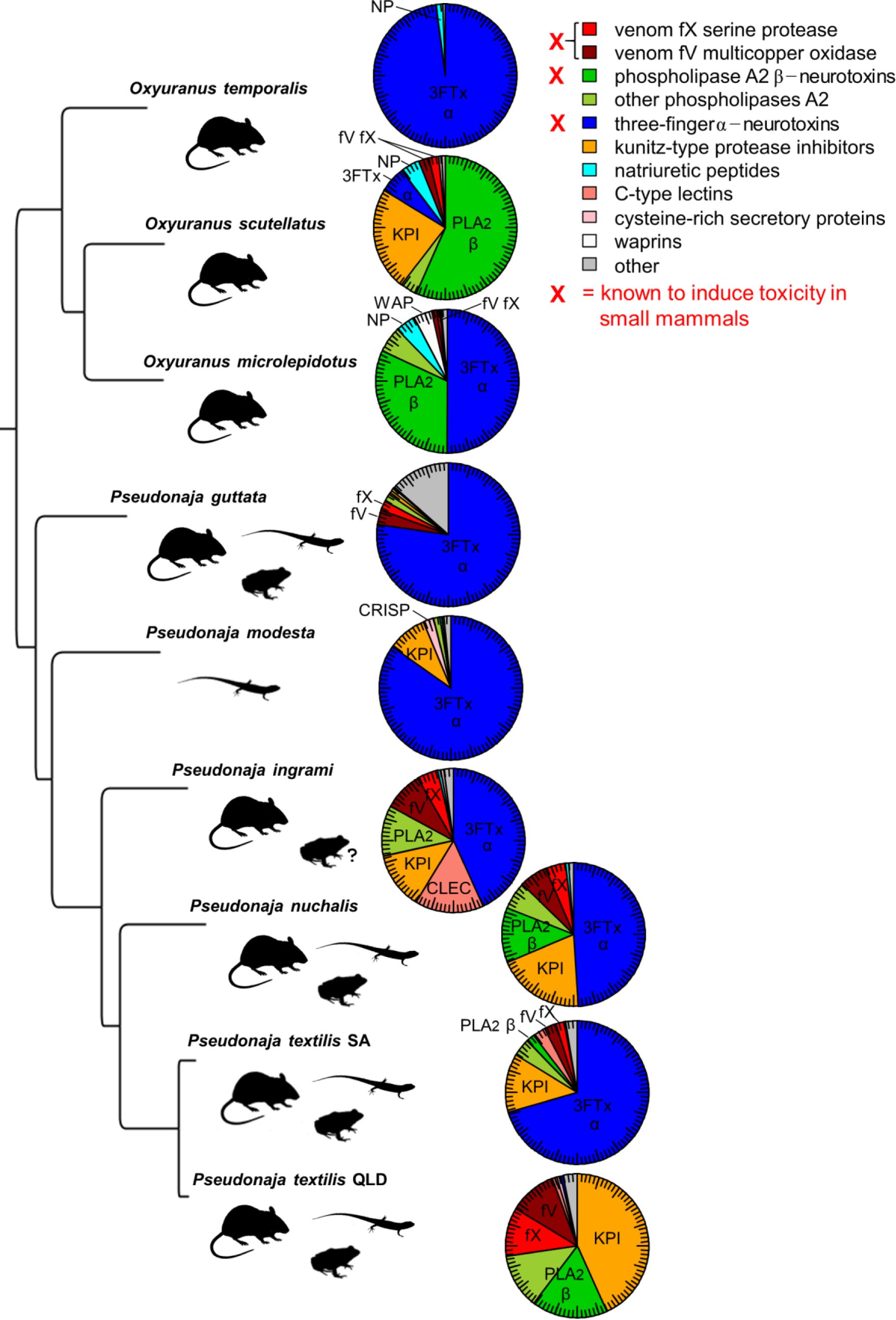
Venom proteome profiles of adult snakes mapped over the species phylogeny. Protein families containing toxins known to induce poisoning in mammals are marked with a red X (whether any of the other protein families found in Australian elapid venoms contain mammal-toxic proteins is currently unknown). Protein expression was estimated using relative iBAQ label-free quantification (major tick = 5%, minor tick = 1% of venom proteome). Main diet items include small mammals, squamate reptiles and amphibians (see Methods).

### Other mammal-eating taipans possess venoms rich in PLA_2_ neurotoxins

Venoms of mammalivorous specialists, the coastal (*Oxyuranus scutellatus*) and inland (*O. microlepidotus*) taipans, contained copious amounts of phospholipase A_2_ presynaptic neurotoxins, i.e. chains belonging to taipoxin and paradoxin clades (Fig. 2, Additional files 1-4). Their relative iBAQ abundances in *O. scutellatus* and *O. microlepidotus* venom samples were 56.6% and 31.8%, respectively. The difference in α-neurotoxin abundance was prominent, *O. microlepidotus* venom containing a considerably greater quantity of these neurotoxins (*O. microlepidotus* iBAQ = 50.1% vs only 5.8% in the *O. scutellatus* sample). There is an indication that α-neurotoxin abundance may vary geographically. For example, a more reduced effect of Saibai Island *Oxyuranus scutellatus* venom on chick biventer cervicis contractile response to acetylcholine suggests that this population would have even lower expression of α-neurotoxins than the specimens from the mainland [39]. *O. scutellatus* and *O. microlepidotus* venoms also contained prothrombinase toxins, but less than in some *Pseudonaja* venoms (Table 1).

### Some brown snakes lack textilotoxin or related PLA_2_ neurotoxins despite feeding on mammals

As with most other *Pseudonaja* species, venoms of Ingram’s brown snake *Pseudonaja ingrami* and the speckled brown snake *P. gutatta* contained prothrombinase toxins and abundant α-neurotoxins (Fig. 2). Interestingly, our samples from these species lacked textilotoxin or related β-neurotoxin homologues (Additional files 1-4). *P. ingrami* feeds on rats, and the lack of potent presynaptic neurotoxins in its venom adds another example of variable expression of mammal-targeting toxins among related species.

### Venoms of several brown snake species contain high levels of prothrombinase toxins

Venom prothrombinase was found in the venoms of all examined *Pseudonaja* that include small mammals in their diet, including *P. textilis, P. nuchalis, P. ingrami* and *P. guttata*. Very high expression levels of venom prothrombinase fX and fV subunits were recorded in the first three mentioned species (Table 1). For instance, in our *P. textilis* venom sample from Queensland (Mackay area), the relative iBAQ proportions were: venom fX = 11.0% and venom fV = 10.6%. A recent study recorded high fX activity of the western brown snake (*P. mendgeni*) venom, suggesting that high expression of prothrombinase toxins can be expected in this venom as well [41].

### Negligible expression of venom prothrombinase in a lizard-eating specialist

Two peptides unique to fX serine protease and four peptides unique to fV were detected in the venom of the ringed brown snake (*P. modesta*) (Additional files 1-3). However, fX and fV expression levels in its venom were extremely low (iBAQ = 0.0035% and 0.0022%, respectively). Their detection indicates that this lizard-eating brown snake apparently has functional genes encoding venom prothrombinase components, but the expression is too low to induce a noticeable procoagulant effect [41, 42]. A transcriptomic study on the West Australian *P. modesta* venom did not find transcripts belonging to fX and recorded low-level expression of fV [38]. Given the down-regulated expression of prothrombinase subunits, there is a possibility that the toxin may be ineffective at subduing squamate reptiles. This is concordant with the lack of prothrombin activation and factor Xa activity by venoms of brown snake juveniles, which also feed on lizards [20, 41].

### Eastern brown snakes from Queensland and South Australia have similar diets but distinct venom proteomes, differing greatly in α-neurotoxin content

Eastern brown snake (*Pseudonaja textilis*) venom from Queensland is characterised by higher abundances of kunitz-type serine protease inhibitors, PLA_2_ neurotoxins (textilotoxin) and venom prothrombinase proteins than South Australian venom of the same species (Fig. 2). The most striking difference in venom composition was high abundance of three finger α-neurotoxins in South Australian *P. textilis* venom (iBAQ = 70.6%) and very low levels of these neurotoxins in Queensland *P. textilis* venom (iBAQ = 0.61%). This finding is of particular interest considering that populations of *P. textilis* from these regions have been reported to have similar diets, eating mainly rodents and lizards [20]. Our previous study found high levels of pseudonajatoxin b in South Australian *P. textilis* venom and very low levels in Queensland *P. textilis* venom [43]. This long-chain α-neurotoxin is lethal to rodents [44], and its down-regulated expression in the venom of rodent-eating Queensland *P. textilis* suggests involvement of factors other than diet influencing venom phenotype. A phylogenetic study of *Pseudonaja* found that coastal Queensland and South Australian *P. textilis* populations belong to different mtDNA lineages [45]. Extensive population-level sampling is necessary to determine the patterns of variation in venom proteome composition within these lineages, which is beyond the scope of the present paper. One study compared venom samples of *Pseudonaja textilis* individuals from South Australian localities and reported similar toxin families but variable expression of protein family members (e.g. textilotoxin subunits) [46].

### Variation in expression of α-neurotoxin types

Alpha-neurotoxin types exhibited different expression patterns in various species (Fig. 3). A maximum likelihood tree of three-finger toxins distinguished two subclades within the oxyuranine α-neurotoxin type, provisionally labelled as A and B (Additional file 4). A striking difference in expression of α-neurotoxin types is seen between a mammal-eating specialist *Oxyuranus temporalis* and a lizard-eating specialist *Pseudonaja modesta*. Venoms of both species were composed predominately of α-neurotoxins, but *O. temporalis* venom contained abundant short- and long-chain α-neurotoxins, whereas that of *P. modesta* was dominated by oxyuranine α-neurotoxins of subclade A (Fig. 3). Hypothetically, certain α-neurotoxins could be adapted to more efficiently target the neuromuscular system of a specific prey class, but this remains unknown. Few α-neurotoxins from Australian elapid venoms have been physiologically characterised, and with these data lacking, it is not possible to say whether any of the observed variation in their expression is caused by prey type selection. It is certain that factors other than prey type selection are involved because of variable expression of the rodent-toxic long-chain α-neurotoxin pseudonajatoxin b and its homologues in rodent-eating species.

**Figure 3.**
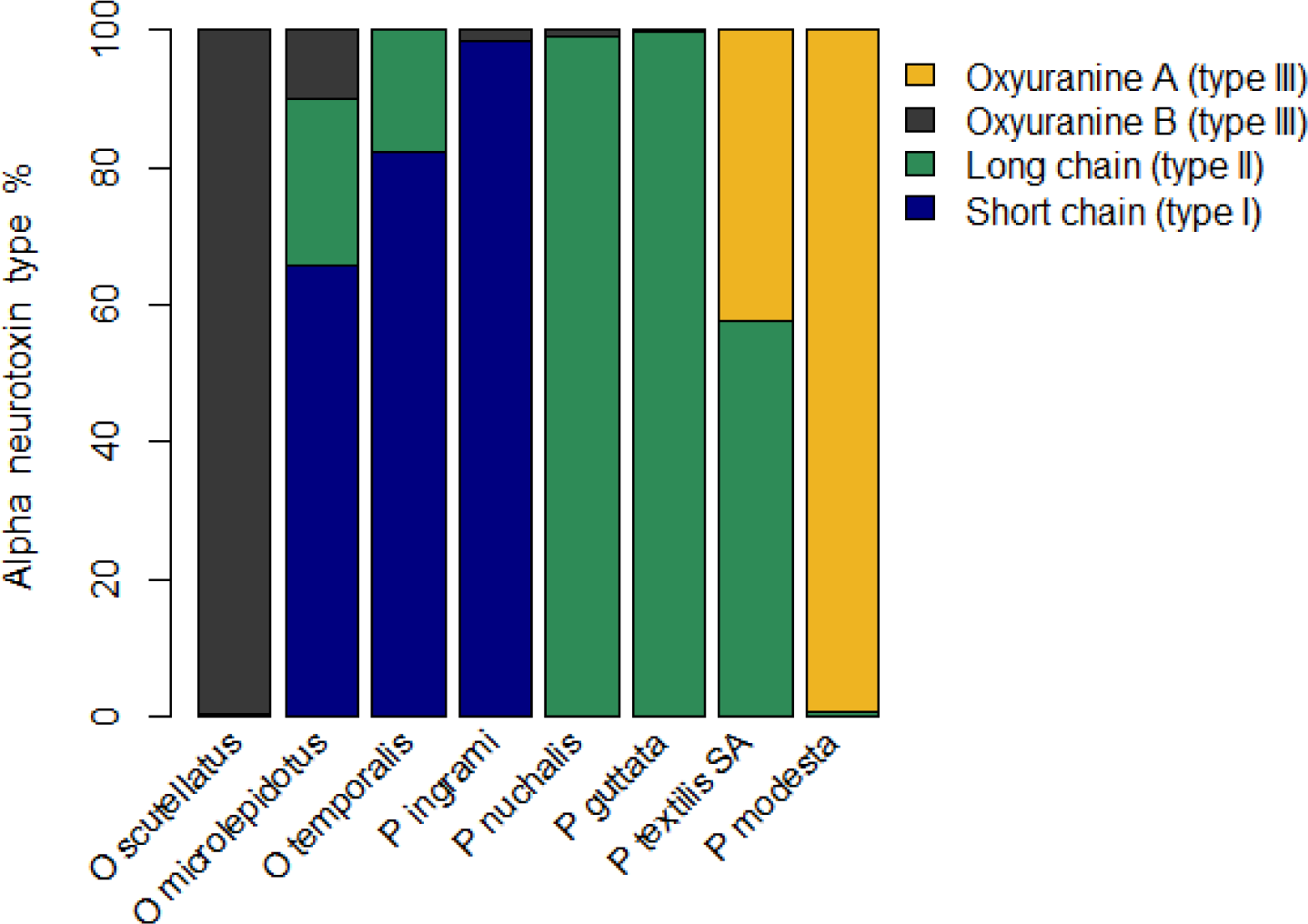
Venoms differed in relative abundances of α-neurotoxin types. See Methods for venom sample details.

### Other main components

Kunitz-type serine protease inhibitors were abundant venom proteome components of most *Pseudonaja* species and *Oxyuranus scutellatus* (Fig. 2). This was by far the most abundant protein family comprising the venom of *Pseudonaja textilis* from Mackay, Queensland (iBAQ = 43.2%). Natriuretic peptides were found in significant quantities in the venoms of all three *Oxyuranus* species, with the iBAQ abundances of around 4% in *O. scutellatus* and *O. microlepidotus*, and a somewhat lower value in *O. temporalis* venom (1.6%). *Pseudonaja ingrami* venom contained high levels of C-type lectin proteins, comprising 15.5% of the expressed proteins. Waprins were found at a significant level in *O. microlepidotus* venom, constituting about 4.2 %. Cysteine-rich secretory proteins (CRISP) had the highest relative expression in the lizard-feeding *Pseudonaja modesta*. Some of these families, such as kunitz-type serine protease inhibitors and natriuretic peptides, possess potent bioactivities with potentially toxic effects in prey [47-50].

### Diverse low-abundance proteins

This analysis identified a myriad of proteins expressed at lower levels (relative iBAQ expression was < 1% in all studied species). Some of these include: nerve growth factor, cystatin, cathepsin, calreticulin, transferrin, 5’nucleotidase, aminopeptidase, snake venom metalloproteinase (SVMP), carboxypeptidase, protein-disulfide isomerase, phospholipase B-like, sulfhydryl oxidase, complement C3, vascular endothelial growth factor (VEGF), hyaluronidase, glutathione peroxidase, PLA_2_ inhibitors, phosphatidylethanolamine-binding protein, alkaline phosphatase, dipeptidase, venom dipeptidyl peptidase and others (Additional files 1-3).

### Venom proteome composition: concluding remarks

Venom prothrombinase is a lethal procoagulant protein complex that induces coagulopathy and circulatory collapse in rodents [29, 51]. Another group of potent toxins, neurotoxic phospholipases A_2_, paralyse the diaphragm in rodent nerve-muscle preparations [52-55].Further, pseudonajatoxin b, a long-chain α-neurotoxin belonging to the three-finger toxin family, is lethal to rodents [44]. It is remarkable that despite the capacity of these toxins to incapacitate small mammals, they show very different expression patterns in closely related mammal-eating Australian elapid snakes. For example, proteomic data suggest that *Oxyuranus temporalis* relies principally on postsynaptic α-neurotoxins to subdue mammal prey, while a related species *Oxyuranus scutellatus* injects a different toxic protein set, containing abundant neurotoxic phospholipases A_2_ and venom prothrombinase.

What processes lead to the observed differences in venom composition? The data presented here and in our previous study that looked at the expression patterns of individual α-neurotoxic proteins in Queensland and South Australian *P. textilis* venoms (see [43]) suggest that up or down-regulation of venom protein expression is important in determining venom proteome composition and associated toxicity. In particular, low-level expression of α-neurotoxins in Queensland *P. textilis* venom indicates that genes coding for these neurotoxins are functional, even though the α-neurotoxin concentration is too low to induce prominent postsynaptic neurotoxicity. Similarly, low-level expression of prothrombinase fX and fV proteins in *Pseudonaja modesta* venom suggests that the species possesses the corresponding genes, but they are expressed at functionally insignificant levels. In addition to protein expression, genomic mechanisms can have a profound impact on venom composition differences in related lineages. For example, a recent study on the evolution of PLA_2_ toxins in rattlesnakes (genus *Crotalus*) suggested that interspecific differences in the content of PLA_2_ toxins are due to gene losses [56]. A role of gene duplication in generating neurotoxin diversity has been demonstrated by a genomic study on *P. textilis*, which identified five paralogous genes encoding oxyuranine α-neurotoxins [57]. Currently, little is known about the contribution of genomic mechanisms and phylogeny to variation in venom composition in Australian elapids.

Diet should intuitively be regarded as an important factor in venom protein evolution because toxins need to efficiently disable physiological functions of targeted prey organisms. However, the results of the current study on *Oxyuranus* and *Pseudonaja* show that prey selection cannot be considered the sole factor contributing to variation in venom proteome composition. This inference is also supported by the finding that the venom of *Bothriechis nigroviridis* (black-speckled palm pitviper) has a unique toxin composition despite the species being observed to eat the same varied vertebrate prey like its congeners, consisting of small rodents, lizards, frogs and small birds [13].

In conclusion, the present study shows that closely related mammal-eating specialist and generalist species of Australian elapid snakes express different sets of venom proteins toxic to mammals. Intraspecific toxin expression patterns and the processes that generate variation in venom composition remain largely unknown. Population-based studies are needed to tackle these questions.

## Methods

### Venoms

Venom samples were obtained from Venom Supplies (Tanunda, SA, Australia) and contained lyophilised crude venom, pooled from adult individuals kept in the facility. The gender was not noted. Adults of *O. temporalis* were housed at Adelaide Zoo (SA, Australia). Details on the venom samples used in this study are given in Table 2.

**Table 2.** Venom samples

| Species / sample | Locality | Number of adults pooled | Supplier |
| --- | --- | --- | --- |
| *Oxyuranus temporalis* | Western Australia (Ilkurlka) | 2 | Adelaide Zoo |
| *Oxyuranus scutellatus* | Queensland (Julatten) | 3 | Venom Supplies |
| *Oxyuranus microlepidotus* | South Australia (Goyders Lagoon) | 6 | Venom Supplies |
| *Pseudonaja guttata* | Queensland |  | Venom Supplies |
| *Pseudonaja modesta* | Queensland | 1 | Venom Supplies |
| *Pseudonaja ingrami* | Queensland (Boulia) | 3 | Venom Supplies |
| *Pseudonaja nuchalis* | Northern Territory (Darwin) | 3 | Venom Supplies |
| *Pseudonaja textilis* | South Australia (Barossa) | 3 | Venom Supplies |
| *Pseudonaja textilis* | Queensland (Mackay) | 2 | Venom Supplies |

### Liquid chromatography and high-resolution mass spectrometry

The venom protein samples were weighed to approximately 1 mg and each sample was reconstituted at a concentration of 5 mg/mL in a solution of 10% acetonitrile, 20 mM ammonium bicarbonate. An aliquot containing 100 µg of protein was reduced in 2.5 mM DTT at 50°C for 15 min followed by alkylation with 10 mM Iodoacetamide for 60 min in the dark at room temperature. Following alkylation, a solution containing 1 µg trypsin (Promega Corp., Madison, WI, USA) in 20 mM ammonium bicarbonate was added and the samples incubated at 37°C overnight. The final concentration of the protein digest mixture was 2 µg/µl.

For analysis by LC-MS, samples were diluted to 0.2 µg/µL using a buffer of 2% acetonitrile and 0.1% formic acid, and centrifuged at 16000 g for 5 min before transfer to an autosampler vial. The mass spectrometer was a Q-Orbitrap, model Q Exactive, Thermo Scientific (Bremen, Germany), software: Thermo Excalibur 2.2. The LC instrument was a nano RS-HPLC, Ultimate 3000, Thermo Scientific (Bremen, Germany), software Thermo Excalibur 2.2 DCMS link.

Tryptic digests were injected in a volume of 2 µL and concentrated on a 100 µm, 2 cm nanoviper pepmap100 trap column with loading buffer (2% acetonitrile, 0.1% formic acid) at a flow rate of 15 µL/min. Peptides were then eluted and separated on a 50 cm Thermo RSLC pepmap100, 75 µm id, 100 Å pore size, reversed-phase nano column with a gradient starting at 97% buffer A (0.1% formic acid), between 4 and 5 min the concentration of buffer B (80% acetonitrile, 0.1% formic acid) increased to 10%, followed by a 30 min gradient to 40% B then to 95% B in 2 min, at a flow rate of 300 nL/min. The eluant was nebulized and ionized using a Thermo nano electrospray source with a distal coated fused silica emitter (New Objective, Woburn, MA, USA) with a capillary voltage of 1800 V.

MS data were acquired with the following parameters: resolution 70000, AGC target 3e6, 120 ms Max IT. Peptides were selected for MSMS analysis in Full MS/dd-MS^2^ (TopN) mode with the following parameter settings: TopN 10, resolution 17500, MSMS AGC target 1e5, 60 ms Max IT, NCE 27 and 3 m/z isolation window. Underfill ratio was at 10% and dynamic exclusion was set to 15 s.

### Proteomic database search

Raw mass spectra were processed with MaxQuant software (version 1.6.0.1), Max Planck Institute of Biochemistry [58-60]. The searched database included 70,087 available protein sequences from UniProt, taxon: Serpentes (downloaded on 17/8/2017). The parameters were: digestion enzyme: trypsin, missed cleavages: 2, fixed modifications: carbamidomethyl (C), variable modifications: oxidation (M), instrument: Orbitrap, first peptide search tolerance: 20 ppm, main peptide search tolerance: 6 ppm, fragment mass tolerance: 0.02 Da. Peptide spectrum matches were filtered with high confidence FDR of 0.01 and protein FDR was set to 0.05.

### Label-free quantification

Venom protein expression was estimated using the iBAQ method, implemented in MaxQuant [61]. iBAQ is calculated by dividing the total intensity of a protein by the number of theoretically observable peptides, produced by *in silico* digestion of proteins with trypsin.

The iBAQ values of individual proteins were summed to obtain the iBAQ value of the whole protein family. This value is then divided by the sum of all iBAQ values to give an estimate of the expression level of a protein family in the venom proteome. The values displayed in Table 1 refer to protein family expression levels in pooled venoms. Expression profiles were graphed using the statistical computing environment R [62].

### Species and toxin trees

For the purpose of plotting diet and venom proteome profiles onto the *Pseudonaja-Oxyuranus* phylogeny, the species tree was constructed using the maximum likelihood method with SMS automatic model selection implemented in PhyML (online version 3.0) [63, 64]. Mitochondrial DNA sequences were available from NCBI GenBank deposited by Skinner et al [45] and Brennan et al [19]. This method produced a tree congruent with the established Bayesian species phylogenies of *Pseudonaja* and *Oxyuranus* [45, 65]. In order to sort identified α-neurotoxins according to their 3FTx subfamily, a maximum likelihood tree was generated using PhyML’s SMS model selection and protein sequences from UniProt. Approximate likelihood ratio test (aLRT) was used for branch support [66]. The same method was applied to assign PLA_2_ matches to textilotoxin, taipoxin and paradoxin neurotoxin clades. Trees were graphed with FigTree software (Andrew Rambaut lab).

### Diet data

Data on diets were obtained from published studies in which prey items were recovered from guts of preserved adult specimens and scats of captured live adults [18-20]. In addition to mammals, *P. ingrami* has also been observed to feed on frogs (N. Dunstan, personal communication).

### Animal ethics statement

Freeze-dried snake venoms used in this study were obtained by the Monash Venom Group, the Department of Pharmacology, Monash University, Australia, in accordance with procedures approved by the Monash University Animal Ethics Committee (MARP/2012/008). Venoms were used for compositional analysis by mass spectrometry, conducted at Monash Biomedical Proteomics Facility, Monash University, Australia.

### Data availability statement

Mass spectrometry proteomics data have been deposited to the ProteomeXchange Consortium via the PRIDE [1] partner repository with the dataset identifier PXD008887.

## Supporting information

## Supplementary Information

Additional file 1: MaxQuant proteinGroups files (xlsx)

Additional file 2: MaxQuant peptide files (xlsx)

Additional file 3: MaxQuant evidence files (xlsx)

Additional file 4: Supplementary figure S1. Phospholipase A_2_ maximum likelihood tree;

Supplementary figure S2. Three-finger toxin maximum likelihood tree (pdf)

## Author Contributions

JS conceived the study, analysed and interpreted the data and wrote the manuscript, DLS designed the LC-MS protocol and performed LC-MS assays, ND conducted venom sampling, contributed expertise and funds, and WCH contributed expertise and funds and reviewed the manuscript. All authors read and approved the final manuscript.

### Acknowledgements

We are very grateful to Terry Morley (Adelaide Zoo, SA, Australia) for provision of *Oxyuranus temporalis* venom. Branka Bruvo-Madaric (Ruder Boskovic Institute, Croatia) kindly provided advice on phylogenetic methods. Many thanks to Brian Bush, Scott Eipper, Stewart Macdonald and Akash Samuel for the permission to use their photographs of *Pseudonaja* and *Oxyuranus* species in the article.

