## Supplementary material for "Venoms of related mammal-eating species of taipans (*Oxyuranus*) and brown snakes (*Pseudonaja*) differ in composition of toxins involved in mammal poisoning"

Supplementary Figures:

Fig. S1. Phospholipase $A_2$ maximum likelihood tree

Fig. S2. Three-finger toxin maximum likelihood tree

### Figure S1

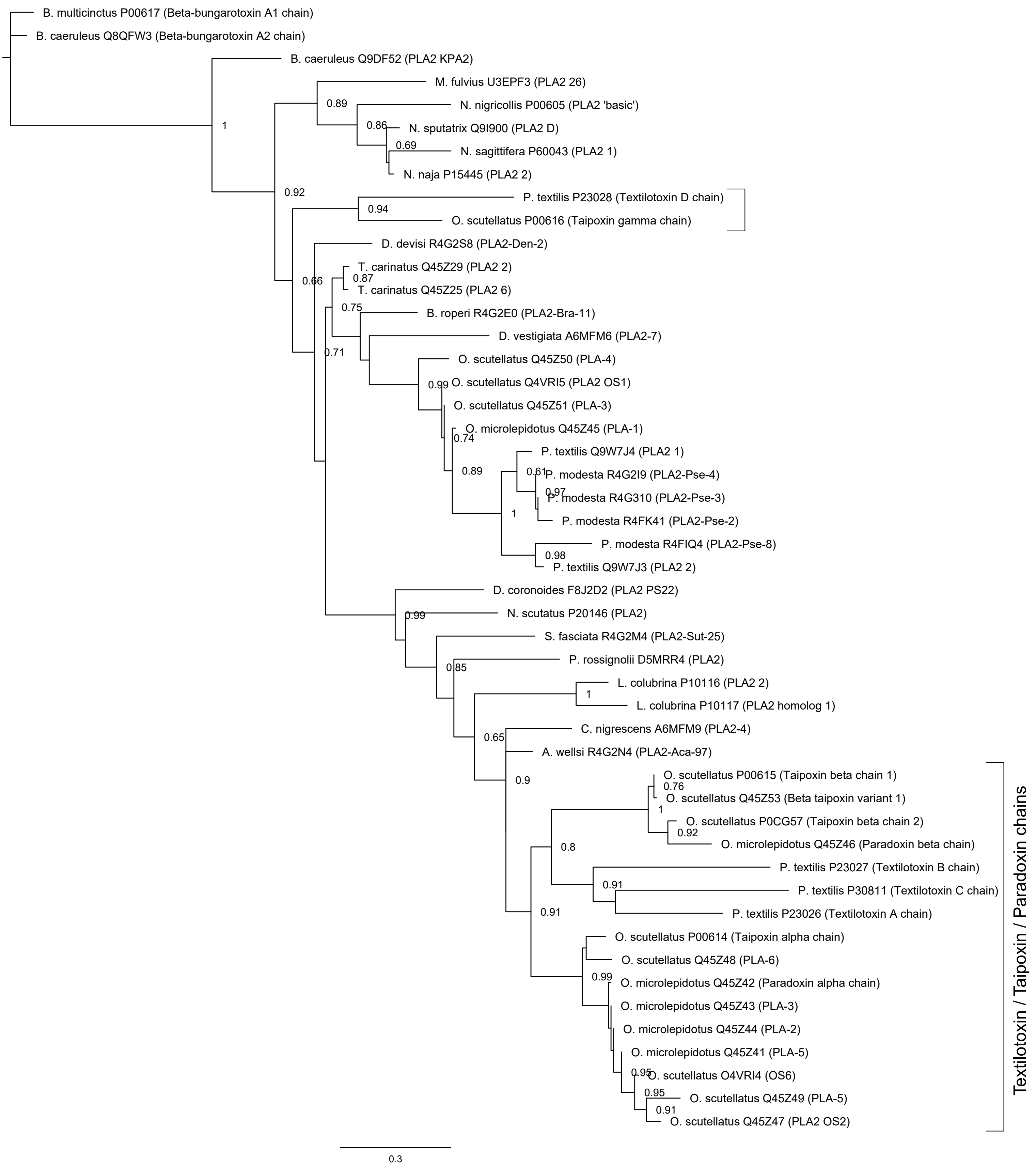
B. multicinctus P00617 (Beta-bungarotoxin A1 chain)
B. caeruleus Q8QFW3 (Beta-bungarotoxin A2 chain)
B. caeruleus Q9DF52 (PLA2 KPA2)
M. fulvius U3EPF3 (PLA2 26)
N. nigricollis P00605 (PLA2 'basic')
N. sputatrix Q9I900 (PLA2 D)
N. sagittifera P60043 (PLA2 1)
N. naja P15445 (PLA2 2)
P. textilis P23028 (Textilotoxin D chain)
O. scutellatus P00616 (Taipoxin gamma chain)
D. devisi R4G2S8 (PLA2-Den-2)
T. carinatus Q45Z29 (PLA2 2)
T. carinatus Q45Z25 (PLA2 6)
B. roperi R4G2E0 (PLA2-Bra-11)
D. vestigiata A6MFM6 (PLA2-7)
O. scutellatus Q45Z50 (PLA-4)
O. scutellatus Q4VRI5 (PLA2 OS1)
O. scutellatus Q45Z51 (PLA-3)
O. microlepidotus Q45Z45 (PLA-1)
P. textilis Q9W7J4 (PLA2 1)
P. modesta R4G2I9 (PLA2-Pse-4)
P. modesta R4G310 (PLA2-Pse-3)
P. modesta R4FK41 (PLA2-Pse-2)
P. modesta R4FIQ4 (PLA2-Pse-8)
P. textilis Q9W7J3 (PLA2 2)
D. coronoides F8J2D2 (PLA2 PS22)
N. scutatus P20146 (PLA2)
S. fasciata R4G2M4 (PLA2-Sut-25)
P. rossignolii D5MRR4 (PLA2)
L. colubrina P10116 (PLA2 2)
L. colubrina P10117 (PLA2 homolog 1)
C. nigrescens A6MFM9 (PLA2-4)
A. wellsi R4G2N4 (PLA2-Aca-97)
O. scutellatus P00615 (Taipoxin beta chain 1)
O. scutellatus Q45Z53 (Beta taipoxin variant 1)
O. scutellatus P0CG57 (Taipoxin beta chain 2)
O. microlepidotus Q45Z46 (Paradoxin beta chain)
P. textilis P23027 (Textilotoxin B chain)
P. textilis P30811 (Textilotoxin C chain)
P. textilis P23026 (Textilotoxin A chain)
O. scutellatus P00614 (Taipoxin alpha chain)
O. scutellatus Q45Z48 (PLA-6)
O. microlepidotus Q45Z42 (Paradoxin alpha chain)
O. microlepidotus Q45Z43 (PLA-3)
O. microlepidotus Q45Z44 (PLA-2)
O. microlepidotus Q45Z41 (PLA-5)
O. scutellatus O4VRI4 (OS6)
O. scutellatus Q45Z49 (PLA-5)
O. scutellatus Q45Z47 (PLA2 OS2)
1
0.89
0.86
0.69
0.92
0.94
0.66
0.87
0.75
0.71
0.99
0.74
0.89
0.61
0.97
1
0.98
0.99
0.85
1
0.65
0.9
0.76
1
0.92
0.8
0.91
0.91
0.99
0.95
0.95
0.91
Textilotoxin / Taipoxin / Paradoxin chains
0.3

Figure S2

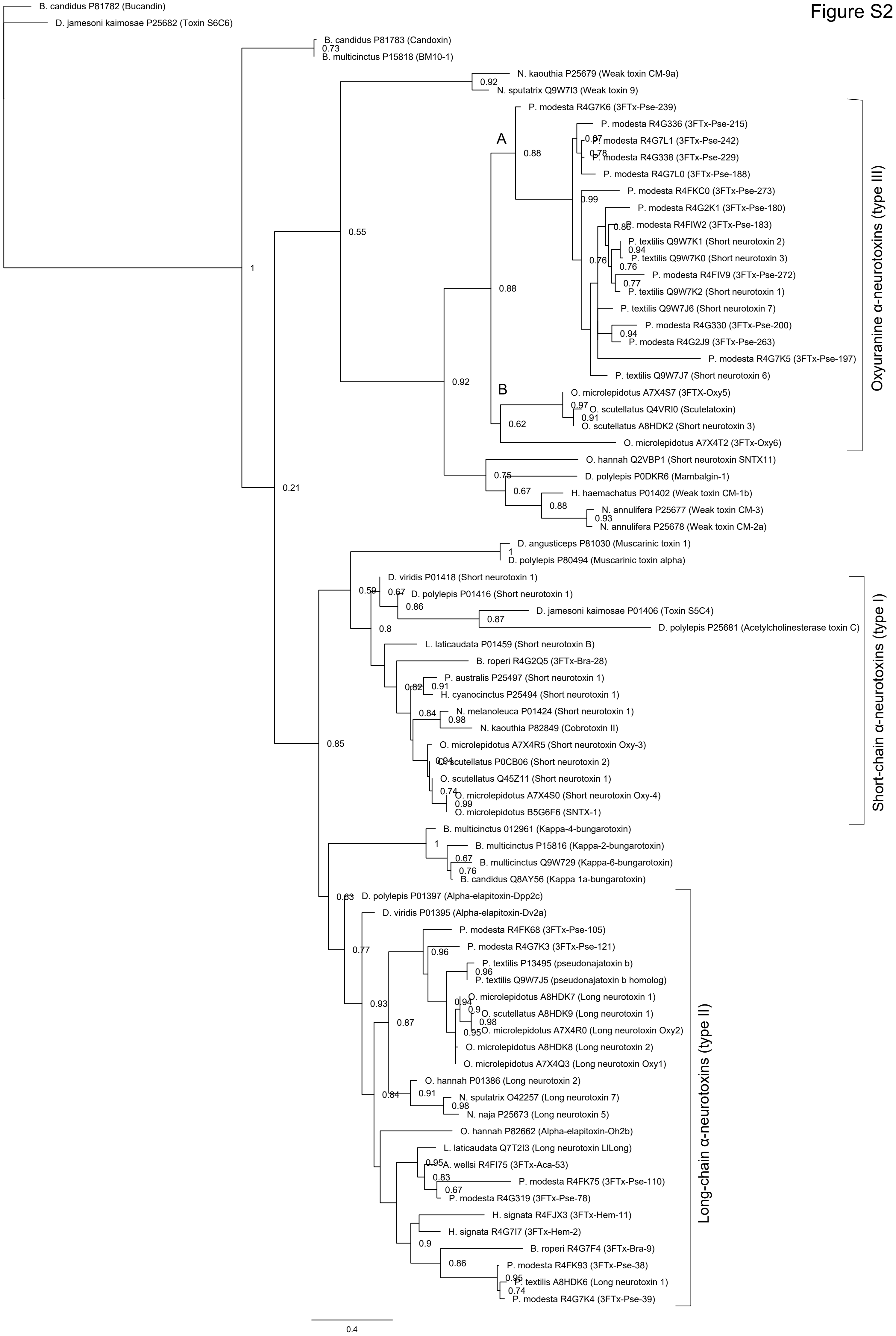
B. candidus P81782 (Bucandin)
D. jamesoni kaimosae P25682 (Toxin S6C6)
B. candidus P81783 (Candoxin)
B. multicinctus P15818 (BM10-1)
N. kaouthia P25679 (Weak toxin CM-9a)
N. sputatrix Q9W7I3 (Weak toxin 9)
P. modesta R4G7K6 (3FTx-Pse-239)
P. modesta R4G336 (3FTx-Pse-215)
P. modesta R4G7L1 (3FTx-Pse-242)
P. modesta R4G338 (3FTx-Pse-229)
P. modesta R4G7L0 (3FTx-Pse-188)
P. modesta R4FKC0 (3FTx-Pse-273)
P. modesta R4G2K1 (3FTx-Pse-180)
P. modesta R4FIW2 (3FTx-Pse-183)
P. textilis Q9W7K1 (Short neurotoxin 2)
P. textilis Q9W7K0 (Short neurotoxin 3)
P. modesta R4FIV9 (3FTx-Pse-272)
P. textilis Q9W7K2 (Short neurotoxin 1)
P. textilis Q9W7J6 (Short neurotoxin 7)
P. modesta R4G330 (3FTx-Pse-200)
P. modesta R4G2J9 (3FTx-Pse-263)
P. modesta R4G7K5 (3FTx-Pse-197)
P. textilis Q9W7J7 (Short neurotoxin 6)
O. microlepidotus A7X4S7 (3FTX-Oxy5)
O. scutellatus Q4VRI0 (Scutelatoxin)
O. scutellatus A8HDK2 (Short neurotoxin 3)
O. microlepidotus A7X4T2 (3FTx-Oxy6)
A
B
Oxyuranine α-neurotoxins (type III)
O. hannah Q2VBP1 (Short neurotoxin SNTX11)
D. polylepis P0DKR6 (Mambalgin-1)
H. haemachatus P01402 (Weak toxin CM-1b)
N. annulifera P25677 (Weak toxin CM-3)
N. annulifera P25678 (Weak toxin CM-2a)
D. angusticeps P81030 (Muscarinic toxin 1)
D. polylepis P80494 (Muscarinic toxin alpha)
D. viridis P01418 (Short neurotoxin 1)
D. polylepis P01416 (Short neurotoxin 1)
D. jamesoni kaimosae P01406 (Toxin S5C4)
D. polylepis P25681 (Acetylcholinesterase toxin C)
L. laticaudata P01459 (Short neurotoxin B)
B. roperi R4G2Q5 (3FTx-Bra-28)
P. australis P25497 (Short neurotoxin 1)
H. cyanocinctus P25494 (Short neurotoxin 1)
N. melanoleuca P01424 (Short neurotoxin 1)
N. kaouthia P82849 (Cobrotoxin II)
O. microlepidotus A7X4R5 (Short neurotoxin Oxy-3)
O. scutellatus P0CB06 (Short neurotoxin 2)
O. scutellatus Q45Z11 (Short neurotoxin 1)
O. microlepidotus A7X4S0 (Short neurotoxin Oxy-4)
O. microlepidotus B5G6F6 (SNTX-1)
Short-chain α-neurotoxins (type I)
B. multicinctus 012961 (Kappa-4-bungarotoxin)
B. multicinctus P15816 (Kappa-2-bungarotoxin)
B. multicinctus Q9W729 (Kappa-6-bungarotoxin)
B. candidus Q8AY56 (Kappa 1a-bungarotoxin)
D. polylepis P01397 (Alpha-elapitoxin-Dpp2c)
D. viridis P01395 (Alpha-elapitoxin-Dv2a)
P. modesta R4FK68 (3FTx-Pse-105)
P. modesta R4G7K3 (3FTx-Pse-121)
P. textilis P13495 (pseudonajatoxin b)
P. textilis Q9W7J5 (pseudonajatoxin b homolog)
O. microlepidotus A8HDK7 (Long neurotoxin 1)
O. scutellatus A8HDK9 (Long neurotoxin 1)
O. microlepidotus A7X4R0 (Long neurotoxin Oxy2)
O. microlepidotus A8HDK8 (Long neurotoxin 2)
O. microlepidotus A7X4Q3 (Long neurotoxin Oxy1)
O. hannah P01386 (Long neurotoxin 2)
N. sputatrix O42257 (Long neurotoxin 7)
N. naja P25673 (Long neurotoxin 5)
O. hannah P82662 (Alpha-elapitoxin-Oh2b)
L. laticaudata Q7T2I3 (Long neurotoxin LlLong)
A. wellsi R4FI75 (3FTx-Aca-53)
P. modesta R4FK75 (3FTx-Pse-110)
P. modesta R4G319 (3FTx-Pse-78)
H. signata R4FJX3 (3FTx-Hem-11)
H. signata R4G7I7 (3FTx-Hem-2)
B. roperi R4G7F4 (3FTx-Bra-9)
P. modesta R4FK93 (3FTx-Pse-38)
P. textilis A8HDK6 (Long neurotoxin 1)
P. modesta R4G7K4 (3FTx-Pse-39)
Long-chain α-neurotoxins (type II)
0.4
